# Characterization and tissue localization of zebrafish homologs of the human *ABCB1* multidrug transporter

**DOI:** 10.1101/2021.02.18.431829

**Authors:** Robert W. Robey, Andrea N. Robinson, Fatima Ali-Rahmani, Lyn M. Huff, Sabrina Lusvarghi, Shahrooz Vahedi, Jordan Hotz, Andrew C. Warner, Donna Butcher, Jennifer Matta, Elijah F. Edmondson, Tobie D. Lee, Jacob S. Roth, Olivia W. Lee, Min Shen, Kandice Tanner, Matthew D. Hall, Suresh V. Ambudkar, Michael M. Gottesman

**Affiliations:** Laboratory of Cell Biology, Center for Cancer Research, National Cancer Institute, National Institutes of Health, Bethesda, MD; Molecular Histopathology Laboratory, Frederick National Laboratory for Cancer Research, Frederick, MD; National Center for Advancing Translational Sciences, National Institutes of Health, Rockville, MD

## Abstract

Given its similarities with mammalian systems, the zebrafish has emerged as a potential model to study the blood-brain barrier (BBB). Capillary endothelial cells at the human BBB express high levels of P-glycoprotein (P-gp, encoded by the *ABCB1* gene) and ABCG2 (encoded by the *ABCG2* gene). However, little information has been available about ATP-binding cassette transporters expressed at the zebrafish BBB. In this study, we focus on the characterization and tissue localization of two genes that are similar to human *ABCB1*, zebrafish *abcb4* and *abcb5*. Cytotoxicity assays with stably-transfected cell lines revealed that zebrafish Abcb5 cannot efficiently transport the substrates doxorubicin and mitoxantrone compared to human P-gp and zebrafish Abcb4. Additionally, zebrafish Abcb5 did not transport the fluorescent probes BODIPY-ethylenediamine or LDS 751, while they were readily transported by Abcb4 and P-gp. A high-throughput screen conducted with 90 human P-gp substrates confirmed that zebrafish Abcb4 has overlapping substrate specificity with P-gp. Basal ATPase activity of zebrafish Abcb4 and Abcb5 was comparable to that of human P-gp. In the brain vasculature, RNAscope probes to detect *abcb4* colocalized with staining by the P-gp antibody C219, while *abcb5* was not detected. Zebrafish *abcb4* also colocalized with claudin-5 expression in brain endothelial cells. Abcb4 and Abcb5 had different tissue localizations in multiple zebrafish tissues, consistent with different functions. The data suggest that zebrafish Abcb4 most closely phenocopies P-gp and that the zebrafish may be a viable model to study the role of the multidrug transporter P-gp at the BBB.

## INTRODUCTION

The human blood-brain barrier (BBB) serves as a defense mechanism for the brain, protecting it from toxins that might gain entry via the blood circulating in the cerebral vasculature [1]. The BBB is comprised of a physical barrier in the form of endothelial cells that form tight junctions to prevent paracellular transport as well as an active barrier comprised of ATP-binding cassette (ABC) transporters expressed on the apical side of endothelial cells to redirect toxins back into the bloodstream [1, 2]. Two transporters that are highly expressed at the BBB are P-glycoprotein (P-gp, encoded by the *ABCB1* gene) and ABCG2 (encoded by the *ABCG2* gene), both of which are associated with multidrug-resistant cancer [3]. The BBB is therefore a significant barrier to the successful treatment of brain malignancies and metastases, as cytotoxic substances, including targeted therapies, are transported by one or both of these transporters [3, 4].

Mouse models have demonstrated the significant and sometimes synergistic role of P-gp and ABCG2 in limiting brain penetration of various compounds. The importance of P-gp at the BBB was demonstrated by Schinkel and colleagues, who discovered serendipitously that mice lacking Abcb1a and Abcb1b (the mouse homologs of human ABCB1) were extremely sensitive to the anthelmintic agent ivermectin [5]. Brain levels of ivermectin were 100-fold higher in knockout mice compared to their wild-type counterparts and mice lacking P-gp were also found to have 3-fold higher levels of vincristine in the brain [5]. The development of mice deficient in both Abcb1a/b as well as Abcg2 demonstrated a role for both transporters in keeping cytotoxins out of the brain. In the case of the mutant BRAF inhibitor encorafenib, mice lacking Abcb1a and Abcb1b demonstrated a 3.4-fold increase in brain levels; mice lacking Abcg2 had 1.8-fold higher levels and mice deficient in Abcb1a/b and Abcg2 had 16.1-fold higher brain levels 2 h after IV administration compared to wild-type mice [6]. The tight junctions formed by the endothelial cells at the BBB limit paracellular transport and augment the effects of transporters such as P-gp and ABCG2, thus resulting in the observed apparent cooperativity of the transporters [7, 8]. This cooperativity has been demonstrated for several small molecule therapies and targeted therapies, including erlotinib and mitoxantrone [3, 8]. While mouse models have provided valuable information with regard to the role of P-gp and ABCG2 at the BBB, they are expensive to maintain and are not amenable to high-throughput screening or direct imaging of the CNS.

The zebrafish (*Danio rerio*) has emerged as a potential model for studying the role of transporters at the BBB [9-11]. Similar to higher vertebrate organisms, zebrafish have a BBB comprised of endothelial cells that form tight junctions characterized by expression of claudin-5 and zona occludens-1 and decreased transcytosis [12-14]. Zebrafish have no single direct homolog of *ABCB1*, but instead express two genes with similar characteristics, *abcb4* and *abcb5* [15, 16]. It has been reported that the C219 antibody that recognizes human P-gp cross-reacts with both zebrafish Abcb4 and Abcb5 [13]. Using this antibody, a protein similar to P-gp was purported to be expressed on endothelial cells that form the zebrafish brain vasculature [9, 13]. However, which of the two proteins (Abcb4 or Abcb5) was expressed at the BBB was unknown. Preliminary studies demonstrated that transporters expressed at the zebrafish BBB are able to efflux known P-gp substrates such as rhodamine 123, loperamide and some tyrosine kinase inhibitors [11, 17]. Both Abcb4 and Abcb5 have been shown to transport some substrates of human P-gp [15, 18, 19]; however, detailed substrate specificity testing of the two transporters has not been performed. Here, we provide a detailed characterization of the zebrafish homologs of human *ABCB1*. Additionally, we localize them to various barrier sites in the fish, suggesting that the zebrafish could indeed be an effective model to study the role of transporters at the human BBB and other sites.

## MATERIALS AND METHODS

### Chemicals

Bisantrene, doxorubicin, mitoxantrone, paclitaxel, rhodamine 123, vinblastine and verapamil were purchased from Sigma-Aldrich (St. Louis, MO). BODIPY FL-ethylenediamine (EDA), BODIPY-prazosin, BODIPY-vinblastine, calcein-AM, and tetramethylrhodamine ethyl ester (TMRE) were obtained from Invitrogen/Life Technologies (Carlsbad, CA). Laser dye styryl 751 (LDS 751) was purchased from Santa Cruz Biotechnology (Dallas, TX). Flutax was from Tocris Bioscience (Minneapolis, MN). AT9283, KW2478 and valspodar were from Apex Biotechnology (Houston, TX). YM-155, VX-680, and 17-AAG were purchased from ChemieTek (Indianapolis, IN). Romidepsin was obtained from Selleck Chem (Houston, TX). Elacridar and tariquidar were from MedChemExpress (Monmouth Junction, NJ).

### Zebrafish husbandry

Animal studies were conducted under protocols approved by the National Cancer Institute and the National Institutes of Health Animal Care and Use Committee. Zebrafish were maintained at 28.5°C on a 14-hour light/10-hour dark cycle according to standard procedures. Larvae were obtained from natural spawning, raised at 28.5°C, and maintained in fish water (60 mg Instant Ocean^©^ sea salt [Instant Ocean, Blacksburg, VA] per liter of DI water). Larvae were checked regularly for normal development. Water was replaced daily. At 5 days post fertilization (dpf), regular feeding was commenced.

### Cell culture

HEK293 cells (ATCC, Manassas, VA) were transfected with empty pcDNA3.1 vector or with vector containing full-length *abcb4* or *abcb5* (all purchased from Genscript, Piscataway, NJ) flanked by a FLAG tag. Sequences were verified before transfection. Transfected cells were grown in MEM medium (Mediatech, Manassas, VA) and were maintained in 2 mg/ml G418 (Mediatech). Clones expressing similar levels of Abcb4 or Abcb5 protein were selected based on FLAG expression as detected by immunoblot. MDR-19 cells that overexpress human P-gp have been previously described [20].

### Flow cytometry

Flow cytometry studies with fluorescent P-gp substrates were performed as previously described [20, 21]. Briefly, trypsinized cells were incubated with the desired fluorescent substrate (150 nM calcein-AM, 5 µM Flutax, 0.5 µM BODIPY-prazosin, 250 nM BODIPY-vinblastine, 0.5 µM BODIPY EDA, 0.5 µM TMRE, 0.5 µM LDS 751, or 0.5 µg/ml rhodamine 123) in the presence or absence of the desired inhibitor (10 µM elacridar, 10 µM tariquidar, 10 µM valspodar, or 100 µM verapamil) for 30 min. Subsequently, the medium was removed and replaced with substrate-free medium in the presence or absence of inhibitor for an additional 1 h. Samples were then analyzed on a FACSCanto II flow cytometer (BD Biosciences San Jose, CA) and data were analyzed with FlowJo software (v 10.6.1, Tree Star, Inc, Ashland, OR).

### High-throughput screen

A high-throughput screen was performed as previously described [22] on empty vector transfected cells, P-gp-overexpressing MDR-19 cells and cells transfected with full-length zebrafish *abcb4* (ZF Abcb4) or *abcb5* (ZF Abcb5). Briefly, cells were plated into 1536-well plates (Corning 7464) at a density of 500 cells/well in 5 µL media. P-gp substrate compounds (90) selected from our previous study [22] were then added at varying concentrations using a 1536-head pin tool (Kalypsis, San Diego, CA) and plates were incubated at 37 °C in 5% CO_2_ for 72 h. CellTiter-Glo reagent (Promega) was dispensed into the wells, incubated for 5 min and luminescence was read on a ViewLux instrument (Perkin-Elmer). Data was normalized based on DMSO basal level (0% activity) and no cell control (-100% activity), which mimics the maximal cell killing effect from this assay. The area-under-the-curve (AUC) was assessed for each compound and was compared for each cell line as previously described [22]. More negative AUC values represented greater cytotoxicity. Compounds were further clustered hierarchically using TIBCO Spotfire 6.0.0 (Spotfire Inc., Cambridge, MA. https://spotfire.tibco.com/) based on AUC values from the screen.

### Immunoblot

Whole cell lysates (30 µg) were obtained from transfected cells using RIPA buffer, heated to 37° for 20 minutes, then subjected to electrophoresis on a premade 4-12% bis-tris gel and transferred to a nitrocellulose membrane. Subsequently the blot was blocked with Odyssey Blocking Buffer (Li-COR Lincoln, NE) for one hour at room temperature, then probed overnight with mouse monoclonal anti-GAPDH (American Research Products, Waltham, MA, 1:8000), the anti-P-gp antibody C219 (Signet Laboratories, Dedham, MA, 1:250), and anti-FLAG antibody (Millipore-Sigma, Milwaukee, WI, 1:1000), where noted. The blot was then incubated with a goat anti-mouse secondary antibody tagged with a near infra-red fluorochrome and fluorescence was measured using the LiCor ODYSSEY CLx (Li-COR).

### Real Time PCR

Total RNA was extracted from HEK cells transfected with empty vector control (Vector), zebrafish *abcb4* (ZF Abcb4), zebrafish *abcb5* (ZF Abcb5), or human *ABCB1*-transfected MDR-19 cells using the RNeasy Mini extraction kit (Qiagen). First strand cDNA synthesis was performed on 500 ng of template RNA using MMLV reverse transcriptase (fc 10 units/µl), dNTP (fc 1mM), Random primer (fc 50 ng/µl) and RNase Inhibitor (fc 1 unit/µl) in 1X First Strand synthesis buffer (Invitrogen/Life Technologies). Gene expression was performed on a BioRad CFX96 real time PCR machine using gene-specific FAM-labelled TaqMan probe assays: human *GAPDH* control ID: Hs99999905_m1(ThermoFisher), zebrafish *abcb4* ID: qDreCIP0043777 and zebrafish *abcb5* ID: 1DreCIP0039260 (both from BioRad). Gene targets (*abcb4* and *abcb5*) were normalized to human *GAPDH* in the HEK-transfected cell lines.

### Cytotoxicity assays

Trypsinized cells were counted and plated into 96-well white opaque plates (5000 cells/well) and allowed to attach overnight. Drugs were subsequently added, with each concentration tested in triplicate, and plates were incubated for 72 h at 37°C in 5% CO_2_. CellTiter-Glo (Promega, Madison, WI) reagent was used to determine cell survival according to the manufacturer’s instructions. Luminescence was subsequently read on a Tecan Infinite M200 Pro microplate reader (Tecan Group, Morrisville, NC). GI_50_ (50% growth inhibitory) concentration values were calculated as the concentration at which 50% luminescence was observed compared to untreated cells.

### ATPase assay

ATPase assays were performed as previously described [23]. Total membranes were isolated from HEK293 cells transfected to express zebrafish *abcb4* (ZF Abcb4), *abcb5* (ZF Abcb5) or human *ABCB1* (MDR-19). Vanadate-sensitive ATPase activity was calculated by measuring the end point phosphate release in the absence and presence of vanadate. Solutions containing 4.0 µg of total membrane protein in 100 µL of ATPase assay buffer (50 mM MES-Tris pH 6.8, 50 mM KCl, 5 mM sodium azide, 1 mM EGTA, 1 mM ouabain, 2 mM DTT, 10 mM MgCl_2_) with 1% DMSO solvent alone (basal activity) or with variable concentrations (0.1, 1 or 10 µM) of the substrates in DMSO were prepared. The tubes were incubated for 3 min at 37°C, after which the reaction was initiated by addition of 5 mM ATP. After a 20-min incubation, the reaction was stopped by addition of 2.5% SDS. The amount of inorganic phosphate released was subsequently quantified by a colorimetric assay [23].

### Immunofluorescence and RNAscope probes

Adult zebrafish were euthanized by submersion in an ice water bath according to an NIH Animal Care and Use Committee-approved protocol, followed by full submersion into fresh 4% paraformaldehyde (PFA) and incubation for 24 h at 4°C. Post-fixation, the fish were submerged in 0.5 M EDTA with a pH of 8.0 and incubated at room temperature for 7 days with gentle agitation. Before processing they were rinsed with 2 washes of nuclease-free PBS. The standard processing protocol for paraffin-embedded samples was used. Two fish were embedded into the same block, with one embedded sagittally and the other coronally. The coronal sections were captured using the bread loafing technique and placed in the block with the tail side down. The tail was removed just past the anus for the coronal sections only. Serial sections from the paraffin blocks were cut at 5 µm and placed on positively charged slides for H&E and multiplex fluorescent images using in situ hybridization (RNAscope) and immunofluorescence. Histopathology was performed by a board-certified veterinary pathologist. Organs evaluated included the following: bone, brain, chromaffin tissue, corpuscle of stannous, esophagus, eye, gall bladder, gill, heart, kidney, hematopoietic tissue, intestine, liver, nares, oral cavity, ovary, pancreas, peripheral nerve, pseudobranch, skeletal muscle, skin, spinal cord, spleen, statoacoustic organ, swim bladder, testis. RNAscope probes for zebrafish abcb4 and abcb5 were purchased from Advanced Cell Diagnostics (Newark, CA). Tissue sections were deparaffinized twice in xylene for 5 minutes each and then twice in 100% ethanol for 3 minutes each. Tissue pretreatment consisted of antigen retrieval in 1X Dako Target Retrieval Solution (pH 9.0) in a RHS-1 Microwave Vacuum Histoprocessor (Milestone Medical, Kalamazoo, MI) at 10 min to 100°C and 25 min at 100°C. Sections were then manually stained with a RNAscope^®^ Multiplex Fluorescent v2 Kit (Advanced Cell Diagnostics) with either a 3-plex Negative control probe, 3-plex positive control probe or with a cocktail of Abcb5-C1 and Abcb4-C2 probes with a 1:750 dilution of TSA-Fluorescein Plus and TSA-Cyanine 3 Plus (PerkinElmer, Shelton, CT), respectively. RNAscope staining was followed by the C219 IHC at a 1:200 dilution for 30 minutes on a Bond RX auto-stainer using the Bond Polymer Refine Kit (Leica, Buffalo Grove, IL) minus DAB & Hematoxylin. Antibody binding was detected with a 1:50 dilution of anti-HRP conjugated with Alexa 594 (Jackson ImmunoResearch Laboratories, Inc., West Grove, PA) for 30 min. Cells were counterstained with 4′,6-diamidino-2-phenylindole (DAPI). Selected organs were evaluated for semi-quantitative scoring for each marker. For colocalization studies with claudin-5, RNAscope staining was followed by claudin-5 (Invitrogen, Grand Rapids, NY, cat # 35-2500) IHC at a 1:50 dilution for 30 minutes on a Bond RX auto-stainer using the Bond Polymer Refine RED Kit (Leica) minus Hematoxylin.

## RESULTS

### Establishment and characterization of HEK293 cells stably expressing zebrafish Abcb4 or Abcb5

In order to characterize the substrate specificity of the zebrafish homologs of human *ABCB1*, we transfected HEK293 cells with either empty vector plasmid, or vector encoding full-length *abcb4* or *abcb5*, with both proteins having a 5’ FLAG tag. Clones were selected based on reactivity with an anti-FLAG antibody, and we chose two clones with similar levels of expression for further study, ZF Abcb4 and ZF Abcb5, as shown in Figure 1A. The predicted molecular weight of zebrafish Abcb4 is approximately 141 kDa and Abcb5 is 147 kDa and the FLAG-tagged proteins ran slightly higher. These clones were also found to have similar levels of gene expression, as measured by PCR analysis (data not shown). The C219 antibody that detects human P-gp has been reported to react with both zebrafish Abcb4 and Abcb5 [13], but this has not been validated with each protein separately. Interestingly, despite the findings with the anti-FLAG antibody, the C219 antibody seemed to preferentially stain Abcb5 over Abcb4 by immunoblot (Figure 1B). The MDR-19 cell line transfected to overexpress human P-gp served as a positive control for C219 staining.

**Fig 1.**
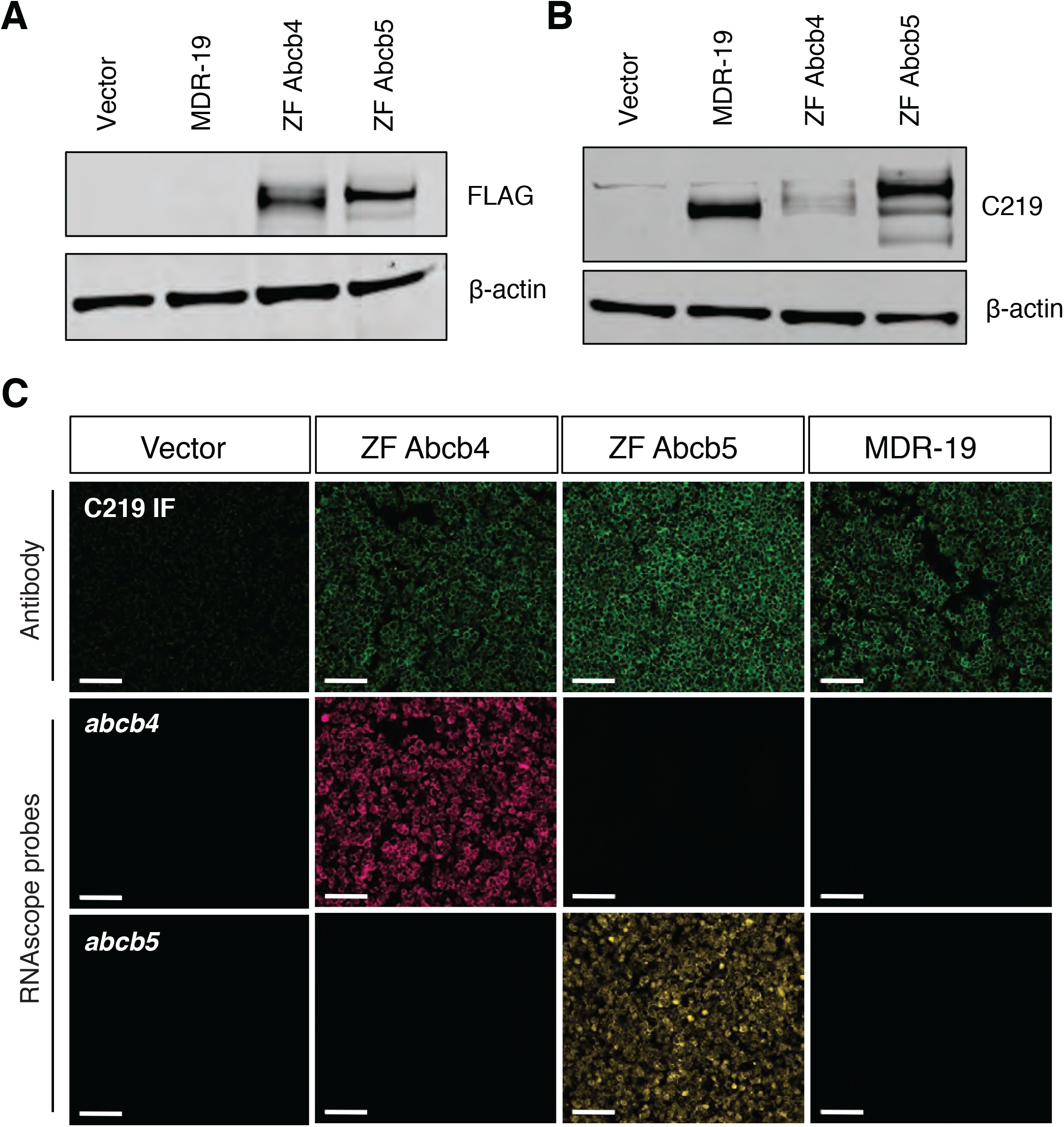
Characterization of cells transfected to express human ABCB1, zebrafish Abcb4 or Abcb5. Whole cell lysates were prepared from HEK293 cells transfected with empty vector (Vector), human ABCB1 (MDR-19), zebrafish abcb4 (ZF Abcb4) or abcb5 (ZF Abcb5), subjected to polyacrylamide gel electrophoresis and transferred to nitrocellulose. Blots were probed with anti-FLAG antibody and beta actin (A) or anti-ABCB1 antibody C219 and beta-actin (B). (C) Paraffin-embedded cells (from A and B) were probed with the C219 antibody (top row, green staining) or RNAscope probes to detect zebrafish *abcb4* (middle row, magenta staining) or zebrafish *abcb5* (bottom row, yellow staining), as outlined in Materials and Methods. Bar = 100 µm.

Transporter expression was measured by immunofluorescence with the C219 antibody and expression of zebrafish Abcb4 and Abcb5 at the cell membrane was noted on both of the selected clones, with slightly more staining of the Abcb5 clone (Figure 1C, top row). The MDR-19 cell line also demonstrated expression of human P-gp (ABCB1) at the cell membrane. The specificity of the RNA *in situ* probes was validated in the transfected cell lines and we found that only the ZF Abcb4 cells stained positively with the *abcb4* probe (Figure 1C, middle row), while only the ZF Abcb5 cells stained positively with the *abcb5* probe (Figure 1C, bottom row); empty vector and MDR-19 cells were not found to react with either probe.

### Cytotoxicity assays demonstrate zebrafish Abcb4 is more similar to human P-gp than zebrafish Abcb5

Having confirmed that our clones all expressed similar levels of zebrafish Abcb4, Abcb5 or human P-gp, we performed 3-day cytotoxicity assays with known substrates of human P-gp: vinblastine, doxorubicin, bisantrene, etoposide, mitoxantrone and paclitaxel. Camptothecin was chosen as a negative control, as P-gp overexpression does not confer resistance to that particular drug. As shown in Figure 2, while both zebrafish Abcb4 and Abcb5 conferred similar levels of resistance as human P-gp to vinblastine and paclitaxel, Abcb4 clearly conferred greater resistance to bisantrene, mitoxantrone, and doxorubicin compared to Abcb5. No cross resistance to camptothecin was observed in any of the cells; GI_50_ values are summarized for the compounds in Table 1. These initial studies suggested that zebrafish Abcb4 and Abcb5 have different substrate specificities and that zebrafish Abcb4 may be functionally similar to human P-gp.

**Fig 2.**
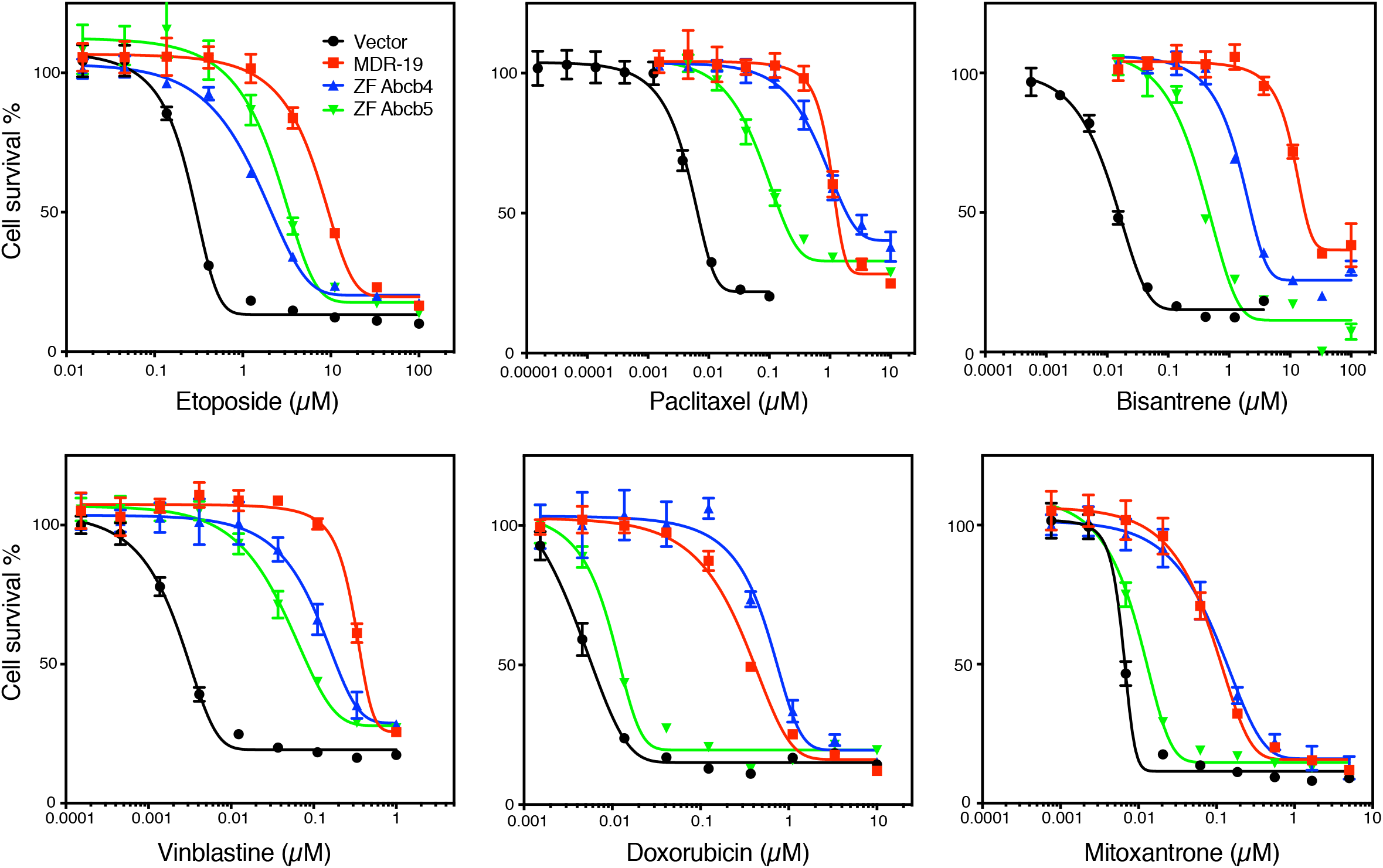
Zebrafish Abcb4 and Abcb5 confer resistance to known P-gp substrates. Three-day cytotoxicity assays were performed with the known human P-gp substrates etoposide, paclitaxel, bisantrene, vinblastine, doxorubicin and mitoxantrone on HEK-293 cells transfected with empty-vector cells (Vector, black curve) or cells expressing human P-gp (MDR-19, red curve), zebrafish Abcb4 (ZF Abcb4, blue curve), or zebrafish Abcb5 (ZF Abcb5, green curve). GI_50_ values were obtained from the curves and are summarized in Table 1.

**Table 1.** Cross-resistance profile of known human P-gp substrates with zebrafish Abcb4 and Abcb5^a^

| Compound | GI<sub>50</sub><br>Vector (μM) | GI<sub>50</sub><br>MDR-19 (μM) | RR*<br>MDR-19 | GI<sub>50</sub><br>ZF Abcb4 (μM) | RR* ZF<br>Abcb4 | GI<sub>50</sub><br>ZF Abcb5 (μM) | RR* ZF<br>Abcb5 |
| --- | --- | --- | --- | --- | --- | --- | --- |
| Bisantrene | 0.016±0.0042 | 18.8±7.2 | 1175 | 1.9±0.56 | 119 | 0.37±0.04 | 23 |
| Camptothecin | 0.009±0.004 | 0.011±0.004 | 1 | 0.009±0.001 | 1 | 0.012±0.004 | 1 |
| Doxorubicin | 0.0092±0.0023 | 0.89±0.63 | 97 | 1.3±0.45 | 141 | 0.021±0.0072 | 2 |
| Etoposide | 0.25±0.062 | 10.0±0.25 | 40 | 2.9±0.62 | 12 | 4.6±1.8 | 18 |
| Mitoxantrone | 0.0053±0.0011 | 0.10±0.017 | 19 | 0.090±0.035 | 17 | 0.010±0.0036 | 2 |
| Paclitaxel | 0.0063±0.0012 | 1.7±0.83 | 270 | 2.5±0.75 | 397 | 0.79±1.05 | 125 |
| Vinblastine | 0.0029±0.0008 | 0.39±0.12 | 134 | 0.26±0.06 | 90 | 0.08±0.01 | 28 |

<sup>a</sup>All compounds were tested at least 3 times. Results presented are mean GI<sub>50</sub> values +/- standard deviation.
\*Relative resistance (RR) value is the ratio of the GI<sub>50</sub> values of MDR-19, ZF Abcb4 or ZF Abcb5 cells to the GI<sub>50</sub> value of empty vector (Vector) cells.

### Efficacy of zebrafish P-gp homolog inhibition is inhibitor- and substrate-dependent

We next compared the ability of human P-gp and zebrafish Abcb4 and Abcb5 to transport known fluorescent substrates of human P-gp using flow cytometry. The substrates tested included calcein AM, rhodamine 123, BODIPY-prazosin, Flutax, BODIPY-vinblastine, LDS 751, BODIPY EDA, and TMRE. We also examined the ability of the human P-gp inhibitors elacridar (10 µM), valspodar (10 µM), tariquidar (10 µM), and verapamil (100 µM) to inhibit transport of the substrates mediated by human P-gp and zebrafish Abcb4 and Abcb5. In the case of MDR-19 cells that overexpress human P-gp (Figure 3A and Supplemental Figure S1), all of the fluorescent probes were transported, as fluorescence of cells incubated with the substrate alone (blue histogram) was lower compared to cells co-incubated with probe and elacridar (orange), tariquidar (light green), valspodar (dark green) or verapamil (pink). Fluorescence histograms of probes co-incubated with inhibitor were overlapping, suggesting complete P-gp inhibition by the various inhibitors.

**Fig 3.**
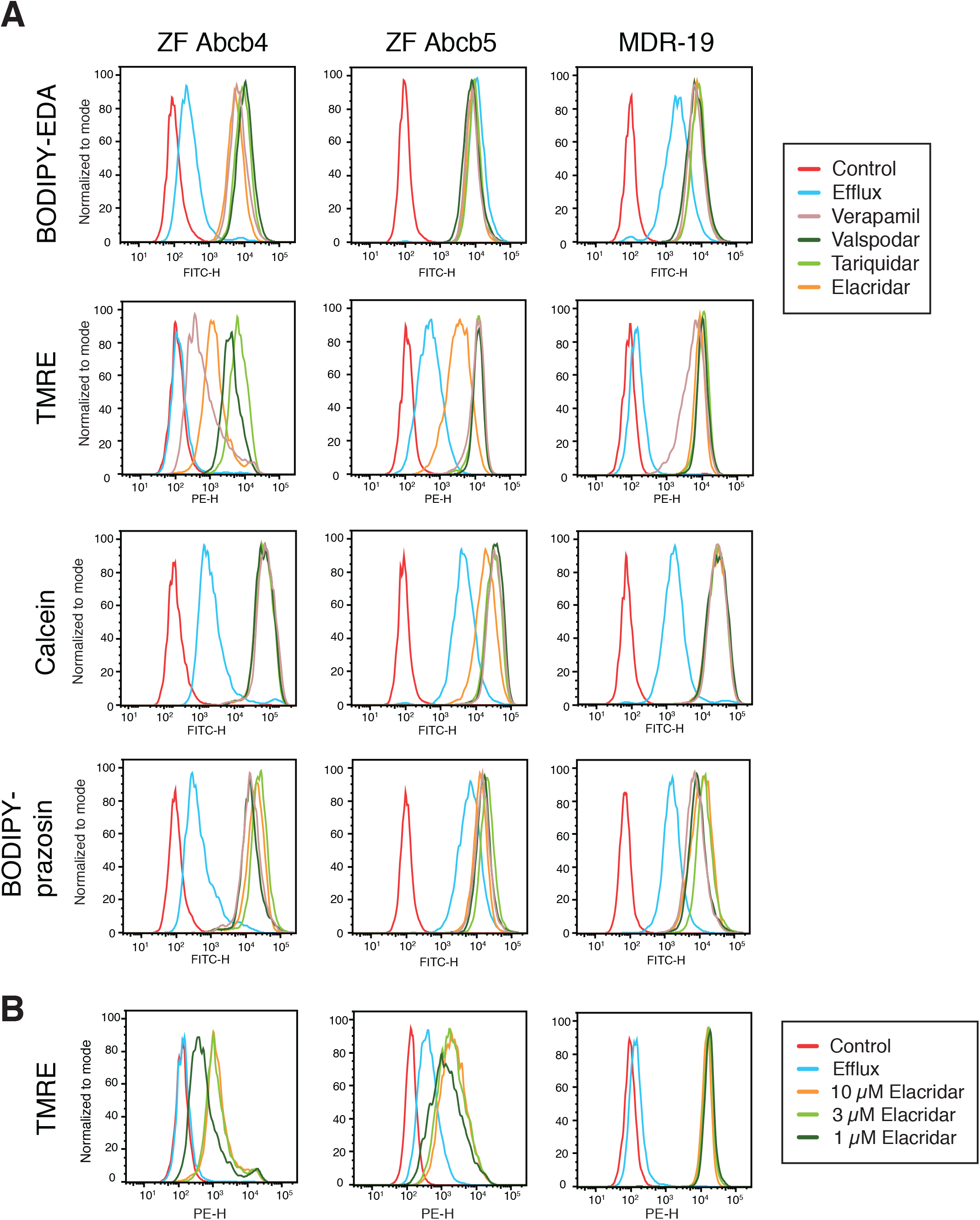
Zebrafish Abcb4 and Abcb5 differentially transport fluorescent P-gp substrates. (A) HEK293 cells transfected to express zebrafish Abcb4 (ZF Abcb4), zebrafish Abcb5 (ZF Abcb5) or human P-gp (MDR-19) were incubated in medium with 0.5 µM BODIPY FL-EDA, 0.5 µM TMRE, 150 nM calcein-AM or 0.5 µM BODIPY prazosin in the presence or absence of 10 µM elacridar, 10 µM tariquidar, 10 µM valspodar, or 100 µM verapamil for 30 min. The medium was removed and replaced with substrate-free medium in the presence or absence of inhibitor for an additional 1 h. (B) Cells from (A) were incubated with medium containing 0.5 µM TMRE in the presence or absence of 1, 3, or 10 µM elacridar for 30 min after which the medium was removed and replaced with substrate-free medium in the presence or absence of inhibitor for an additional 1 h.

We found that zebrafish transporter substrates could be grouped into three categories. First, BODIPY-EDA (Figure 3A, top row) and LDS 751 (Supplemental Figure S1, top row) were transported by Abcb4, but not Abcb5. The intracellular fluorescence of these substrates was decreased in cells expressing Abcb4, as shown by the blue histogram, whereas this histogram is closely aligned with the inhibitor histograms in the case of the Abcb5-overexpressing cells, signifying no transport. Second, although some substrates were transported by both Abcb4 and Abcb5, human P-gp inhibitors did not fully inhibit the zebrafish transporters. This was the case for TMRE (Figure 3A, second row) and rhodamine 123 (Supplemental Figure S1, second row). In the case of Abcb4 and TMRE, both the verapamil (pink) and elacridar (orange) histograms appeared to the left of the tariquidar (light green) and valspodar (dark green) histograms, denoting incomplete inhibition of transport. A similar result was observed with Abcb5 and TMRE, although only elacridar (orange) appeared to the left of histograms for the other inhibitors. Elacridar, verapamil and tariquidar were less effective than valspodar at inhibiting Abcb4-mediated rhodamine transport, while elacridar was less effective at inhibiting Abcb5-mediated rhodamine transport. The other substrates—calcein AM and BODIPY-prazosin (Figure 3A third and fourth row) as well as Flutax, and BODIPY-vinblastine (Supplemental Figure S1, third and fourth row)—fell into the third category, where all appeared to be transported and all inhibitors seemed to be equally effective at inhibiting the zebrafish transporters.

To validate the results observed with elacridar, we performed a dose response, using TMRE as the substrate. In the case of P-gp, elacridar at the lowest concentration of 1 µM (dark green histogram) completely inhibited P-gp-mediated TMRE efflux (Figure 3B). In the case of zebrafish Abcb4 or Abcb5, even 10 µM was unable to completely inhibit TMRE transport. Thus, while the zebrafish transporters Abcb4 and Abcb5 are similar to P-gp, there are some differences.

### High throughput screening of known P-gp substrates

We recently described the development of a high throughput screen to identify substrates of human P-gp [22]. Here, we compared the substrate specificity of zebrafish Abcb4 and Abcb5 to human P-gp by calculating area under the curve (AUC) values for dose-response curves generated from the transfected cells that were tested with of a panel of 90 cytotoxic human P-gp substrates [22]. Results from the screen are shown in the heat map in Figure 4A. Unsupervised clustering demonstrated that Abcb4 was functionally similar to human P-gp. As seen in Figure 4B, the AUC values for zebrafish Abcb4 correlated better with human P-gp (r = 0.94) than with Abcb5 (r = 0.67). Data for each of the cell lines tested with the panel of 90 cytotoxic human P-gp substrates was deposited in PubChem with AID 1508636 for MDR-19 cells (https://pubchem.ncbi.nlm.nih.gov/bioassay/1508636), 1508637 for empty vector cells (https://pubchem.ncbi.nlm.nih.gov/bioassay/1508637), 1508635 for ZF Abcb4 cells (https://pubchem.ncbi.nlm.nih.gov/bioassay/1508635) and 1508633 for ZF Abcb4 cells (https://pubchem.ncbi.nlm.nih.gov/bioassay/1508633).

**Fig 4.**
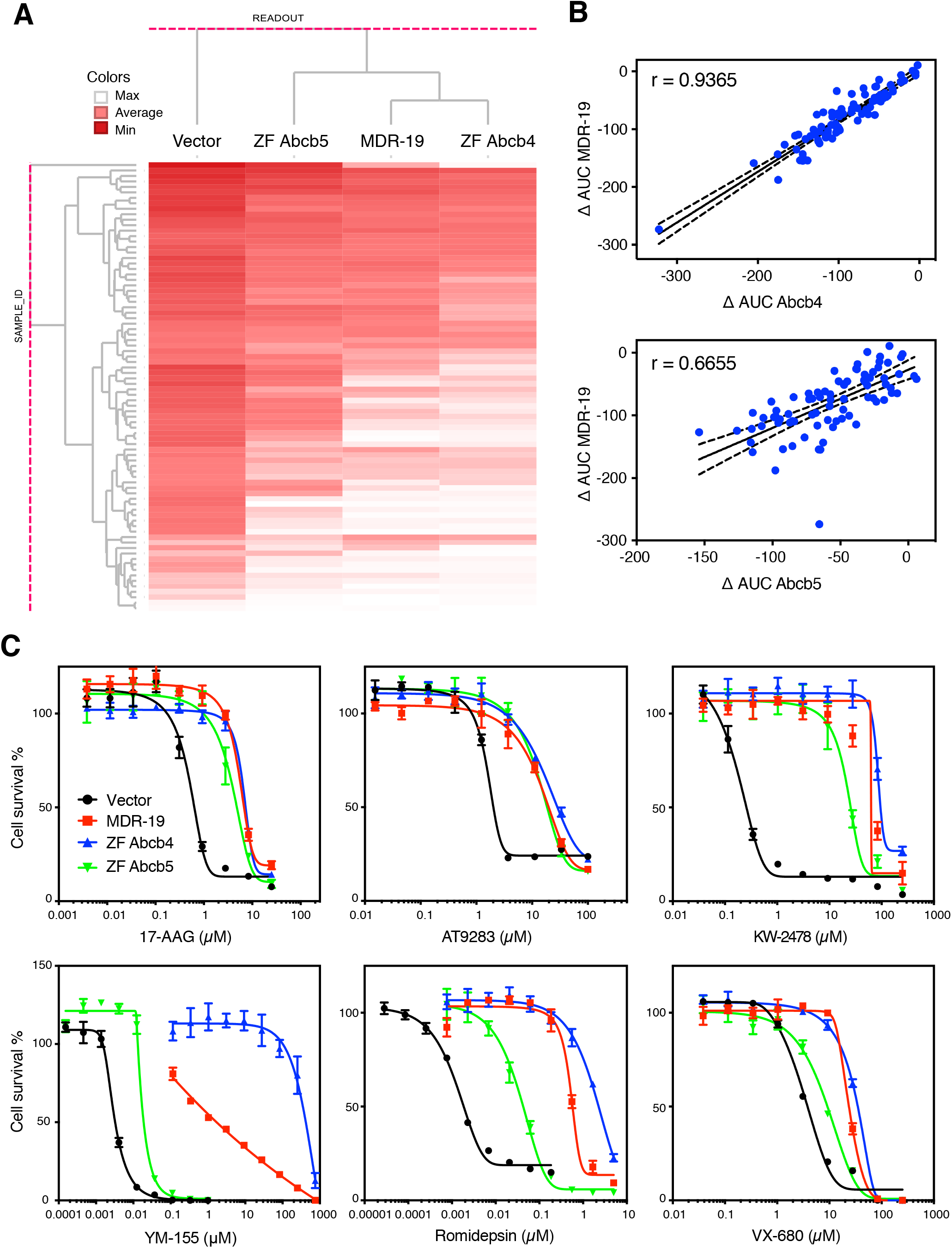
High-throughput screening demonstrates that substrate specificity of zebrafish Abcb4 closely resembles that of human P-gp. (A) High throughput screening was performed with 90 human P-gp substrates with empty vector transfected cells (Vector) or cells transfected to express zebrafish Abcb4 (ZF Abcb4), zebrafish Abcb5 (ZF Abcb5) or human P-gp (MDR-19). Compound activity was then subjected to unsupervised clustering against Vector, ZF Abcb4, ZF Abcb5 and MDR-19 cells, where deep red represents greatest cytotoxicity. (B) The difference in AUC (delta AUC) was determined as the difference between Vector cells as compared to ZF Abcb4, ZF Abcb5 or MDR-19 cells. The delta AUC values for human P-gp (MDR-19 cells) were compared to those of zebrafish Abcb4 (ZF Abcb4) or zebrafish Abcb5 (ZF Abcb5). Solid line represents line of best fit and dashed lines are 95% confidence intervals. (C) Three-day cytotoxicity assays were performed with 6 substrates included in the screen—17-AAG, AT9283, KW-2478, romidepsin, VX-680 and YM-155. Results from all of the experiments are summarized in Table 2.

Of the 90 substrates that were tested, 6 were subsequently selected to confirm the results of the high-throughput screen. We performed confirmatory three-day cytotoxicity assays with 17-AAG (HSP90 inhibitor), AT9283 (Janus kinase/aurora kinase inhibitor), KW-2478 (HSP90 inhibitor), YM-155 (survivin inhibitor), romidepsin (histone deacetylase inhibitor), and VX-680 (tozasertib, aurora kinase inhibitor), on empty vector transfected cells or cells transfected to express zebrafish Abcb4, Abcb5 or human P-gp (data shown in Figure 4C). Comparing the cytotoxicity profiles of the 6 substrates in the transporter-expressing HEK cells, we found that resistance to 17-AAG, AT9283 and KW-2478 was largely similar in the three transporter-expressing cell lines. However, for some compounds, Abcb4 was clearly the better transporter, most notably for YM-155, as zebrafish Abcb5 conferred only low levels of resistance to that compound compared to zebrafish Abcb4 or human P-gp. Data for the 6 compounds are summarized in Table 2. We conclude that, in terms of substrate specificity, zebrafish Abcb4 is more like human P-gp and that zebrafish Abcb5, while functional, confers lower levels of resistance to some substrates.

**Table 2.** Cross-resistance profile of selected compounds from high-throughput screening^a^

| Compound | GI<sub>50</sub><br>Vector (μM) | GI<sub>50</sub><br>MDR-19 (μM) | RR*<br>MDR-19 | GI<sub>50</sub><br>ZF Abcb4 (μM) | RR* ZF<br>Abcb4 | GI<sub>50</sub><br>ZF Abcb5 (μM) | RR* ZF<br>Abcb5 |
| --- | --- | --- | --- | --- | --- | --- | --- |
| 17-AAG | 0.53±0.35 | 4.5±2.4 | 8 | 7.0±0.82 | 13 | 3.8±0.66 | 7 |
| AT9283 | 1.9±0.26 | 14.0±5.3 | 7 | 19.3±1.2 | 10 | 20.7±8.3 | 11 |
| KW-2478 | 0.44±0.31 | 88.3±53.5 | 200 | 140±26 | 318 | 37±27 | 84 |
| Romidepsin | 0.0026±0.0012 | 0.85±0.56 | 327 | 2.3±0.31 | 884 | 0.053±0.023 | 20 |
| VX-680 | 3.0±0.71 | 14.8±7.5 | 5 | 36.5±2.9 | 12 | 8.1±1.8 | 3 |
| YM-155 | 0.0027±0.00059 | 6.2±4.8 | 2296 | 375±66 | 138888 | 0.021±0.0059 | 8 |

<sup>a</sup>All compounds were tested at least 3 times. Results presented are mean GI<sub>50</sub> values +/- standard deviation.
\*Relative resistance (RR) value is the ratio of the GI<sub>50</sub> values of MDR-19, ZF Abcb4 or ZF Abcb5 cells to the GI<sub>50</sub> value of empty vector (Vector) cells.

**Table 3.** Expression pattern of zebrafish *abcb4* and *abcb5*

| Organ | *abcb4* | *abcb5* |
| --- | --- | --- |
| Brain | +++ | - |
| Liver | +++ | + |
| Kidney* | ++ | ++ |
| Skin | - | +++ |
| Ovary | + | +++ |
| Intestine | +++ | - |
| Gill | - | +++ |

Selected organs were evaluated for semi-quantitative scoring for each marker. \*Distinct regions of the nephron are positive for either *abcb4* or *abcb5* (not both).

### ATPase activity of zebrafish homologs of human ABCB1

We next compared the ATPase activity of human P-gp to zebrafish Abcb4 and Abcb5. As shown in Figure 5A, basal levels of ATPase activity were similar among P-gp, zebrafish Abcb4 and Abcb5. For all proteins, ATPase activity was stimulated by verapamil and basal ATPase activity could be inhibited by the addition of 1 µm tariquidar (Figure 5B). This is in agreement with a previous report in which verapamil-inducible ATPase activity was demonstrated for zebrafish Abcb4 [15]. Furthermore, we demonstrate that inhibition of Abcb4 activity by tariquidar is similar to that observed with human P-gp.

**Fig 5.**
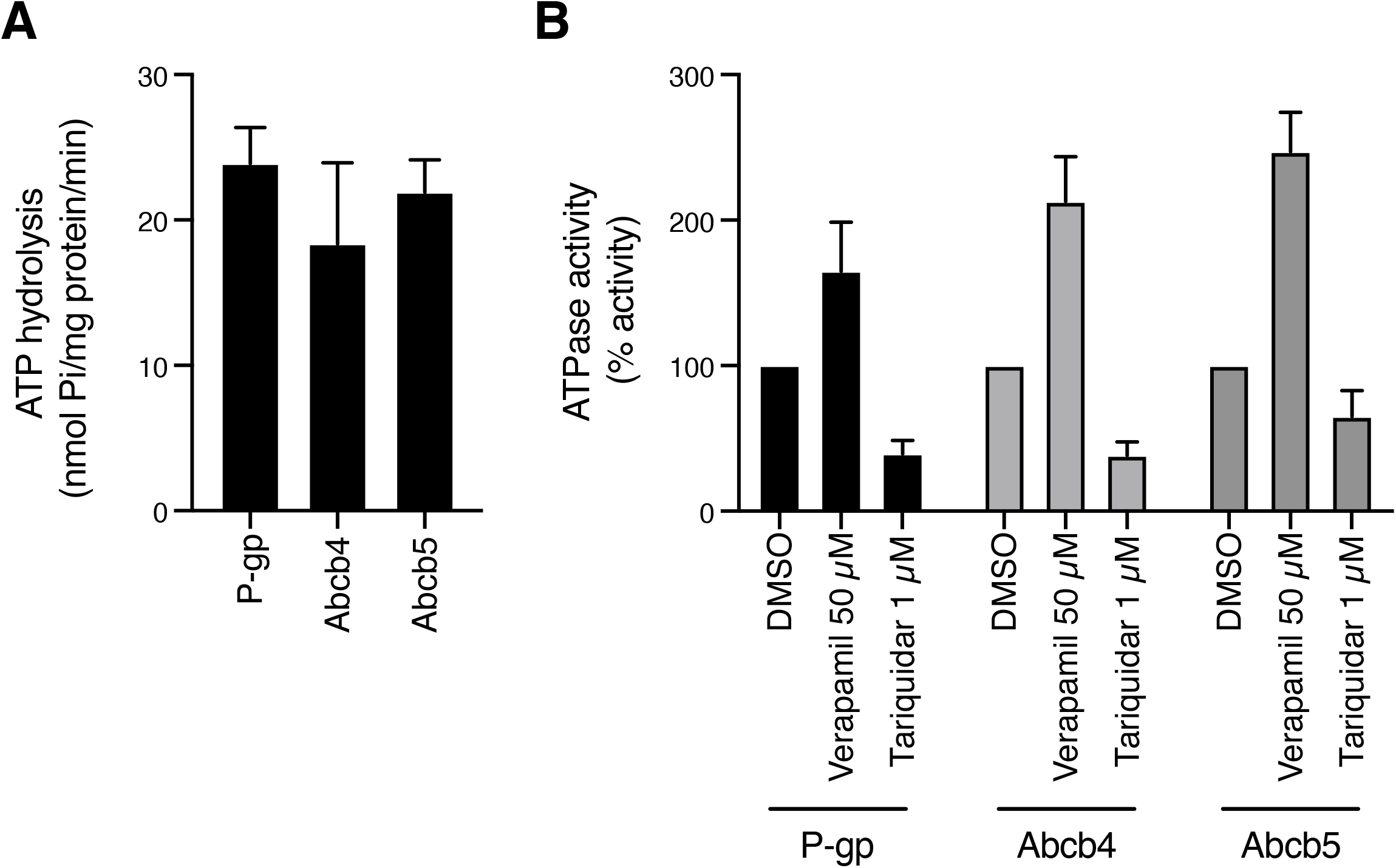
ATPase activity of zebrafish Abcb4 and Abcb5 is similar to that of human P-gp. (A) The vanadate-sensitive activity of zebrafish Abcb4, zebrafish Abcb5 and human P-gp was determined as outlined in Materials and Methods using membranes isolated from ZF Abcb4, ZF Abcb5 or MDR-19 cells, respectively. (B) Basal P-gp ATPase activity (DMSO) was compared to activity in the presence of 50 µM verapamil, or in the presence of 1 µM tariquidar. Graphs from (A) and (B) depict average values from three independent experiments (error bars +/-SD).

### Molecular modeling of zebrafish Abcb4 and Abcb5 reveals potential differences in the drug-binding pocket

To look for potential explanations for the differences in substrate and inhibitor specificity between P-gp and zebrafish Abcb4 and Abcb5, homology models of each protein were generated using the Swiss-model program (https://swissmodel.expasy.org/). The human P-gp structure PDB:6QEX was used as a template, which corresponds to P-gp in complex with taxol and the Fab of the UIC2 antibody [24]. The Clustal Omega program (https://www.ebi.ac.uk/Tools/msa/clustalo/) was used to align the sequences relative to human P-gp. Figure 6A shows the binding pocket of human P-gp with taxol bound at the center of the cavity. The model is split into two halves to provide a clearer view of both sides of the molecule. We found 64% and 58% amino acid identity between P-gp and zebrafish Abcb4 and Abcb5, respectively. The differences between the amino acids in the transmembrane region between P-gp and zebrafish Abcb4 (Figure 6B) and Abcb5 (Figure 6C) were then interrogated. Amino acids were color-coded based on their similarity to the corresponding amino acid in human P-gp with color progressing from green (most similar) to yellow (fairly similar) to red (least similar). Both proteins showed differences with human P-gp in the drug binding pocket; however, Abcb5 had more significantly different amino acids relative to P-gp, as shown by increased red colored residues in the drug-binding pocket. These structural differences in the residues potentially explain the difference in substrate specificity between the proteins.

**Fig 6.**
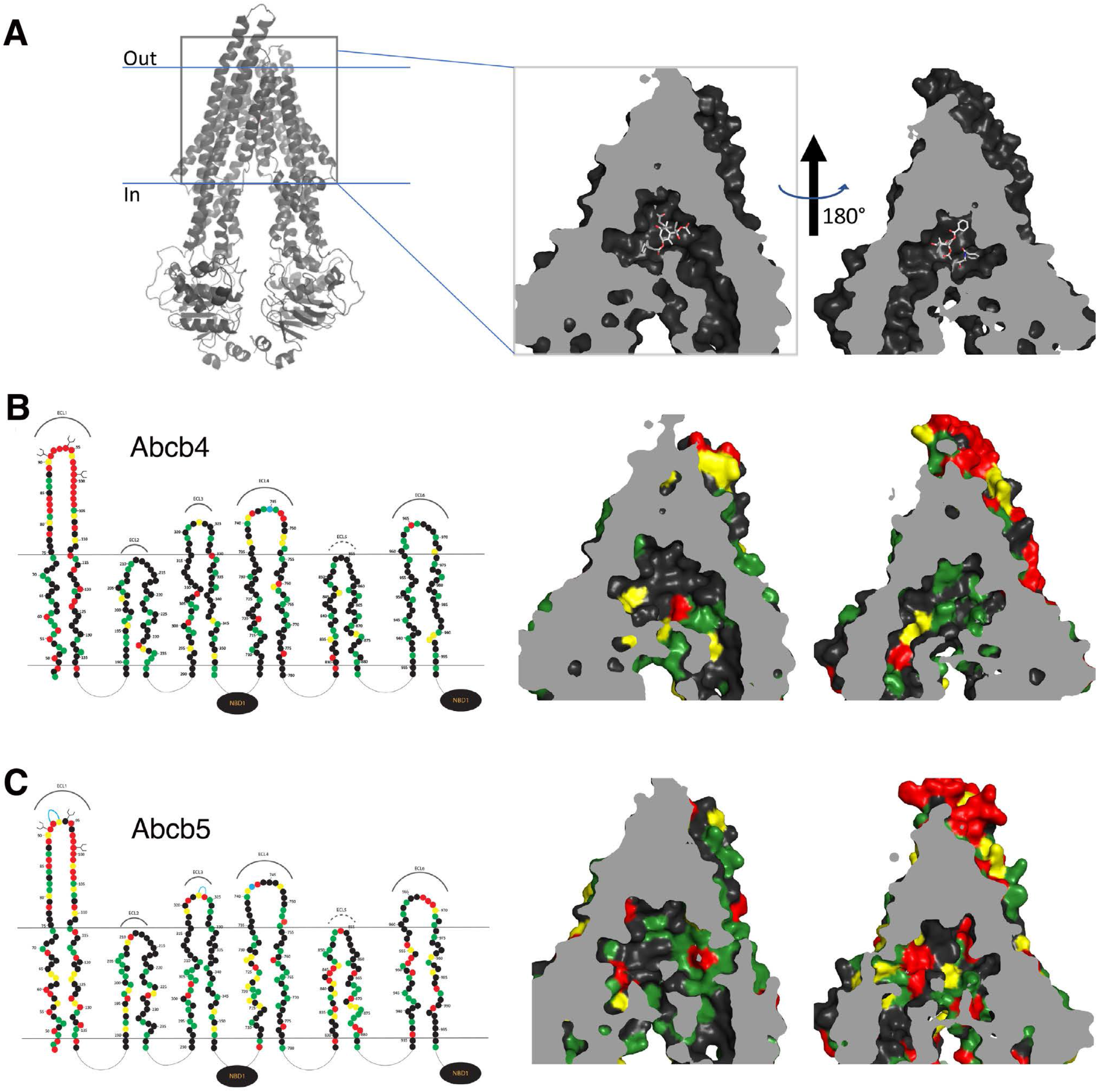
3D homology modeling of amino acid similarity in the binding pocket of zebrafish Abcb4 and Abcb5. (A) 3D modeling of the human P-gp structure that models P-gp in complex with taxol and the Fab of the UIC2 antibody (PDB:6QEX). The model is split into two halves to provide a clearer view. Amino acid differences in the transmembrane regions of the models of human P-gp and zebrafish Abcb4 (B) and Abcb5 (C) are shown. In (B) and (C), on the left side linear 2D models of Abcb4 and Abcb5 are shown. Amino acids (filled circles) similar to human P-gp are shown in green (most similar), yellow (fairly similar) or red (least similar).

### Immunohistochemical localization of Abcb4 and Abcb5 in adult zebrafish

In light of the different substrate specificities of the two different homologs, we decided that it was important to localize the transporters in the zebrafish to determine whether the zebrafish could be used as a model to study the human blood-brain barrier. We first performed immunohistochemistry analysis of the zebrafish P-gp isoforms using the C219 antibody. Staining of the whole fish is shown in Figure 7A, with the negative control shown in Figure 7B. We found C219 reactivity in the vasculature of the brain, including the cerebellum, mesencephalon and telencephalon (Figure 7C). Higher magnification of the cerebellum revealed staining consistent with brain vasculature (Figure 7D). Staining was also noted in the gill epithelium (Figure 7E), a subset of renal tubules (Figure 7F), enterocytes of the gastrointestinal tract, hepatocytes, and ovarian follicles (data not shown). This expression pattern is similar to that of humans, where P-gp is expressed in the BBB, the GI tract, liver, and the kidneys [25].

**Fig 7.**
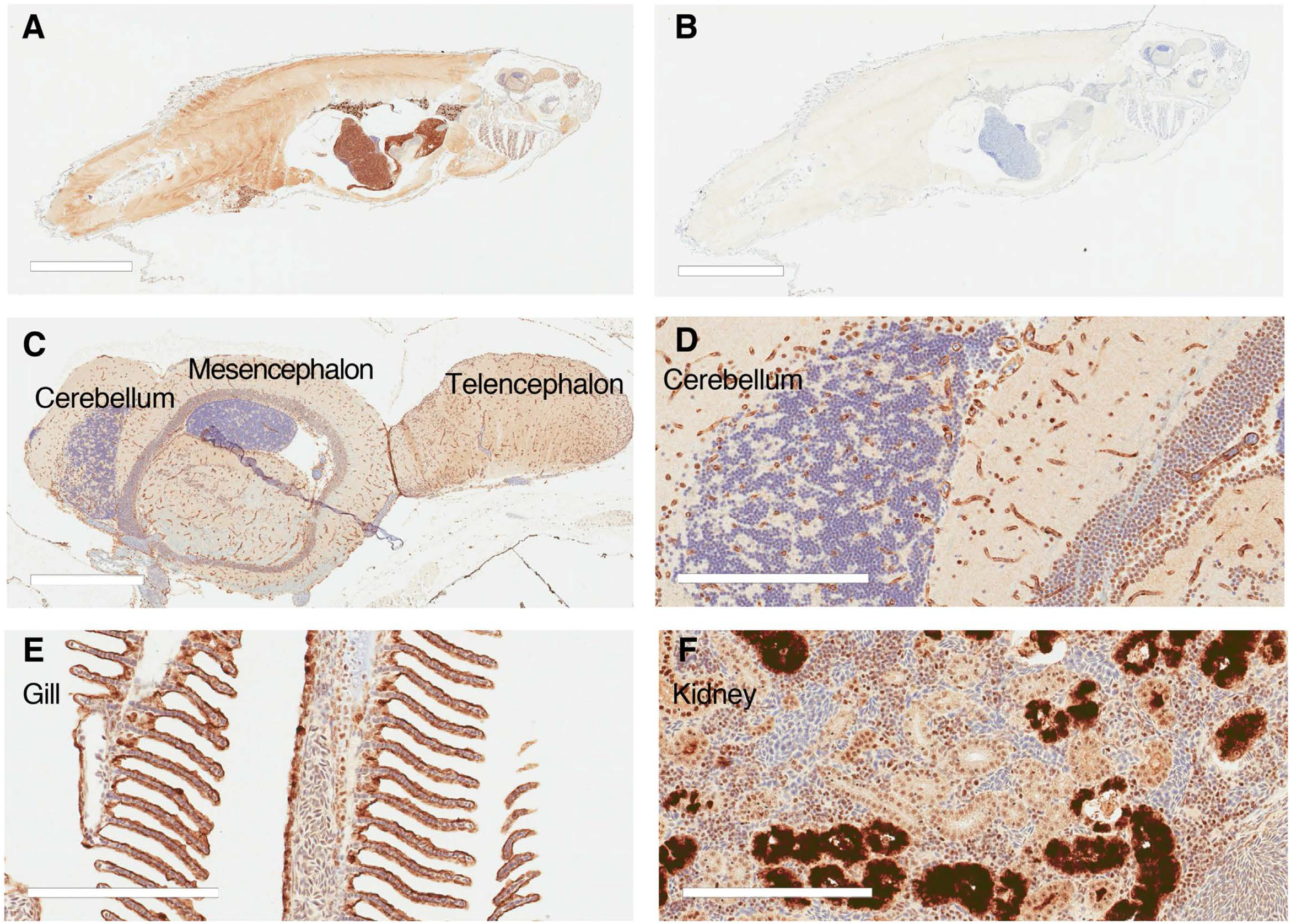
Immunohistochemical staining of Abcb4 and Abcb5 with the C219 antibody in adult zebrafish. Whole adult zebrafish were stained with the C219 antibody (A) or negative control (B), as noted in the Materials and Methods. Bar = 5 mm for (A) and (B). C219 staining of the zebrafish brain (C) with an enlarged portion of the cerebellum (D). Positive staining of C219 antibody was also noted in the gill (E) and kidney (F). Bar = 600 µm for (C) and 200 µm for (D-F).

### Organ-specific expression patterns are observed for zebrafish abcb4 and abcb5

As the C219 antibody cannot qualitatively differentiate between the two zebrafish isoforms, as demonstrated in Figure 1C, we coupled antibody staining with RNAscope technology that recognizes RNA expression. In the brain microvasculature, we found that the C219 antibody (magenta) colocalized with the probe for *abcb4* (yellow), while the *abcb5* probe (green) was not detected (Figure 8, left column). Similar results were observed in the gastrointestinal tract, where *abcb4* (yellow) was colocalized with the C219 antibody (magenta), but *abcb5* (green) was not detected (Figure 8, right column). High levels of C219 staining (magenta) were observed in the renal tubules of the kidney and distinct regions of the nephron stained positive for either *abcb4* (yellow) or *abcb5* (green, Figure 9A). Interestingly, high levels of predominantly *abcb5* were observed in ovarian follicles (Figure 9B). The expression patterns of abcb4 and abcb5 in various organs are summarized in Table 2.

**Fig 8.**
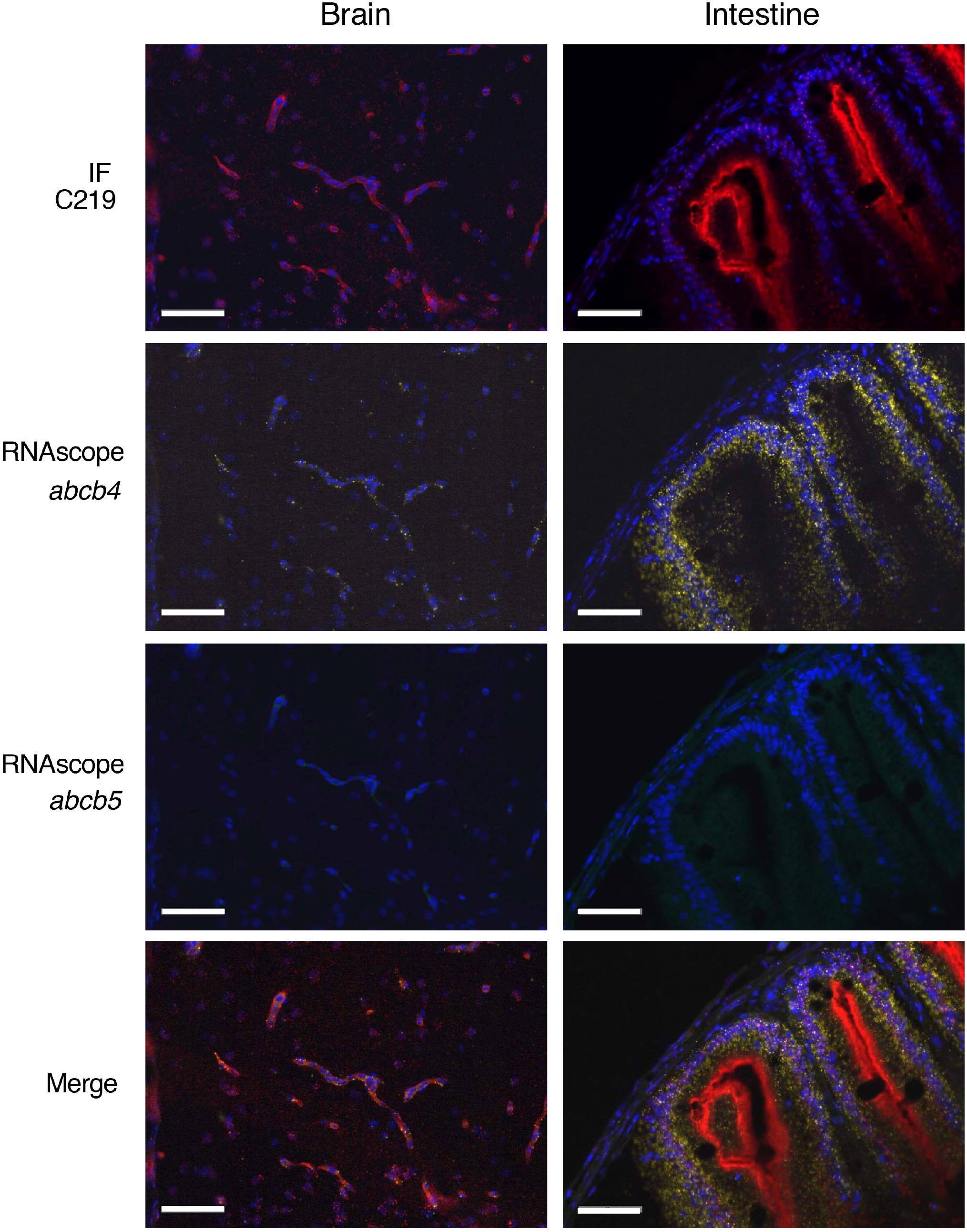
Zebrafish *abcb4* colocalizes with C219 staining in the zebrafish brain and gastrointestinal tract of adult zebrafish. Zebrafish sections were stained with RNAscope^®^ *abcb4*-C2 (yellow) and *abcb5*-C1 (green) probes followed by the C219 antibody (red) as outlined in Materials and Methods. Fluorescence channels were interrogated individually and merged in sections of the zebrafish brain (left column) or gastrointestinal tract (right column). Bars = 50 µm. Nuclei were stained with DAPI (blue).

**Fig 9.**
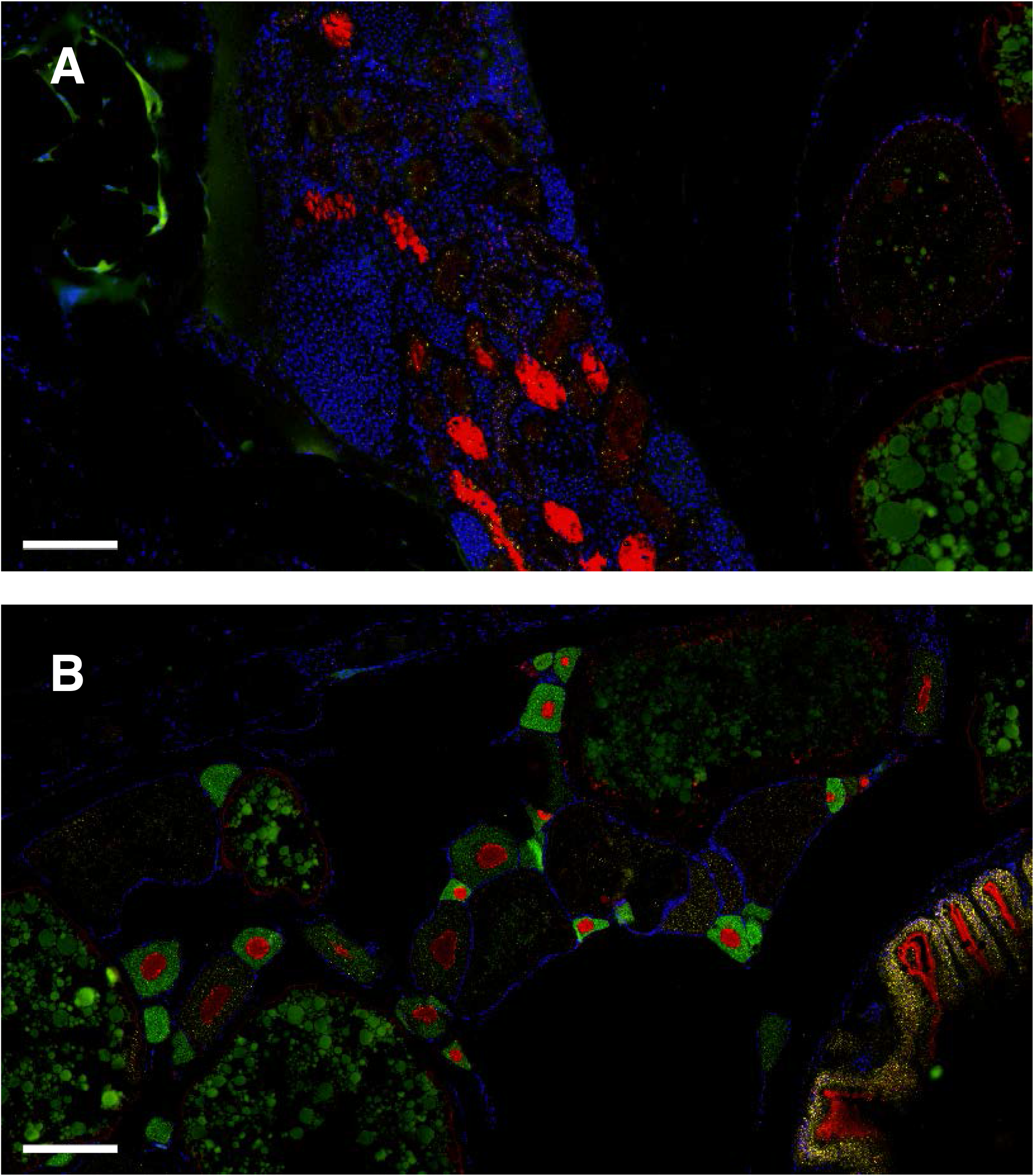
Colocalization of zebrafish *abcb4* and *abcb5* with C219 staining in the zebrafish kidney and ovary. Zebrafish sections were stained with RNAscope^®^ *abcb4*-C2 (yellow) and *abcb5*-C1 (green) probes followed by the C219 antibody (red) as in Figure 8. Nuclei were stained with DAPI (blue). Fluorescence channels were merged in sections of the zebrafish kidney (A) or ovary (B). Bar = 200 µm.

To confirm Abcb4 expression in the brain vasculature, we stained for the tight junction protein claudin-5 (magenta) in the mesencephalon and combined this with the *abcb4* probe (green, Figure 10). The *abcb4* probe clearly colocalized with claudin-5. Zebrafish Abcb4, with a substrate specificity that phenocopies human P-gp, appears to be the only *ABCB1* isoform in the brain vasculature of the zebrafish.

**Fig 10.**
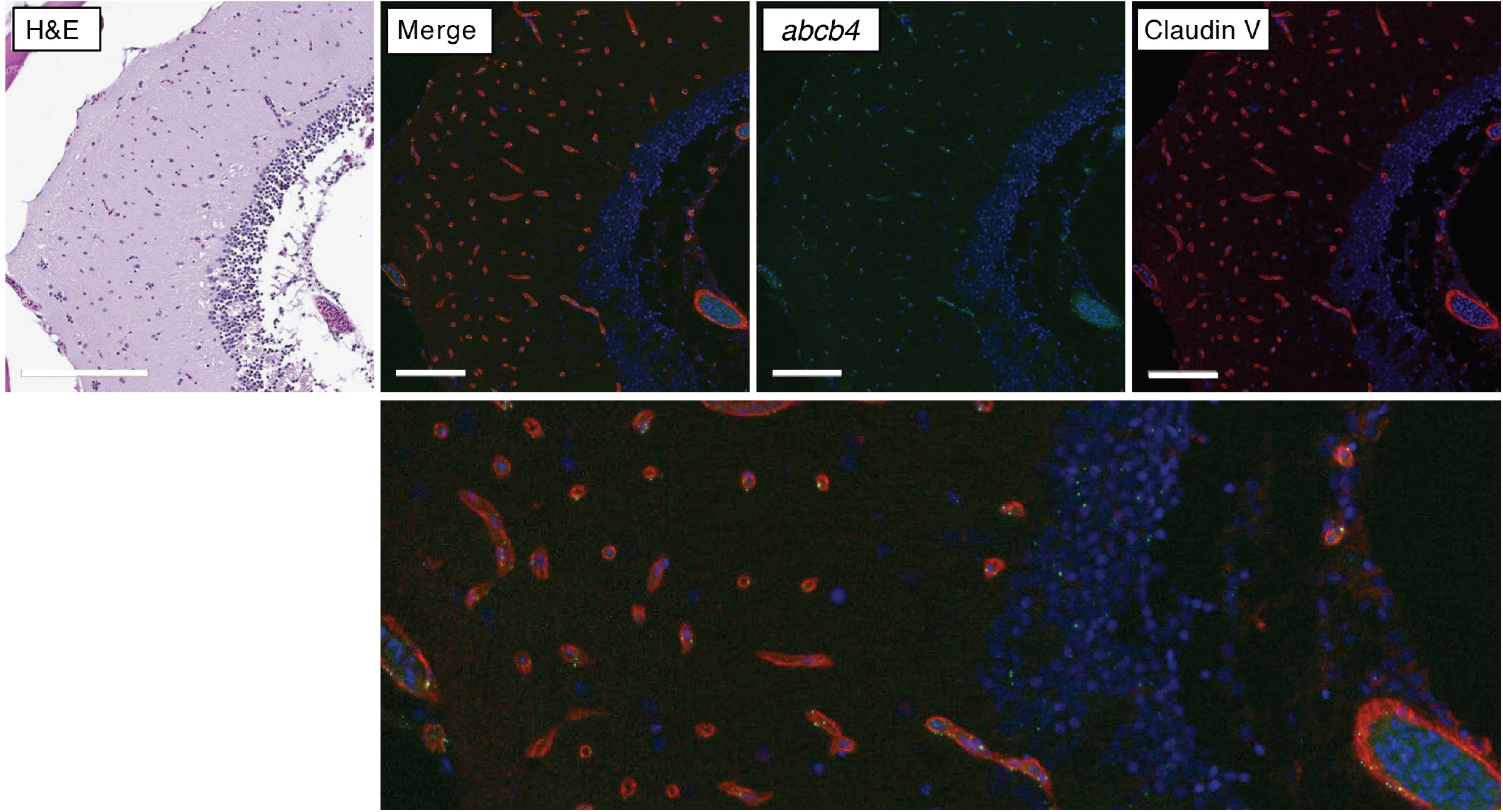
Zebrafish *abcb4* colocalizes with claudin-5 in the zebrafish brain. Zebrafish sections were stained with H&E, or RNAscope^®^ *abcb4*-C2 (green) followed by an antibody to claudin-5 (red) as outlined in Materials and Methods. Nuclei were stained with DAPI (blue). Fluorescence channels were interrogated individually and merged. Lower figure is a higher magnification view of the merged image. Bar = 200 µm for H&E image and 100 µm for upper fluorescence images.

## DISCUSSION

The zebrafish is emerging as a suitable model for studying the human BBB, and with the development of oral dosing techniques, it may also be useful in modeling the oral bioavailability of drugs [26]. While mammals are known to express ABC transporters at the BBB and in the gastrointestinal tract, the localization and substrate specificity of homologous transporters in the zebrafish had not been carefully studied. Here, we report that zebrafish Abcb4 is the predominant transporter expressed at the zebrafish BBB and intestine, that both transporters are expressed in various regions of the kidney, and that Abcb5 is nearly exclusively expressed in the gill, skin and ovary. Additionally, in terms of substrate specificity, zebrafish Abcb4 is functionally similar to human P-gp, suggesting that zebrafish *abcb4* is actually the homolog to human *ABCB1*. Based on transporter localization and specificity, the zebrafish may represent a valuable model for studying the oral bioavailability of compounds and the uptake of drugs into the brain.

Like the mammalian BBB, the endothelial cells of the zebrafish BBB form tight junctions characterized by expression of ZO-1 and claudin-5 and are in close contact with pericytes [27]. However, zebrafish appear to lack homologous astrocytic foot processes at the BBB and instead have radial glia cells that do not sheathe the endothelial cells to the extent observed in mammals, leaving their function at the BBB unclear [27, 28]. While previous reports have suggested that a homolog of human P-gp is expressed at the zebrafish BBB, this was concluded based on reactivity with the anti-human P-gp antibody C219, which was purported to recognize both zebrafish homologs of *ABCB1* [9, 13]. Consistent with these previous reports, we also found C219 reactivity in the endothelial cells of the zebrafish brain, and using RNA in situ techniques, have identified *abcb4* as the gene responsible for the observed C219 staining. Thus, the zebrafish could indeed be a powerful tool to study the role of transporters in limiting the brain penetration of drugs, and initial studies have shown that the zebrafish could be comparable to mouse models to determine whether drugs can cross the BBB [11].

Concerning the two zebrafish proteins that are similar to human P-gp, we find that both zebrafish Abcb4 and Abcb5 transport known P-gp substrates, but that Abcb5 has a slightly narrower substrate specificity profile compared to Abcb4. Previous work by Fischer and colleagues found that Abcb4 could transport several fluorescent P-gp substrates, but they suggested that Abcb5 might not be a xenobiotic transporter based on morpholino knockdown of *abcb5* in zebrafish embryos [15]. More recently, Gordon et al. reported efflux of fluorescent P-gp substrates in zebrafish ionocytes and found this to be due to overexpression of *abcb5* [18]. Our work expands on these previous reports, as we show that both zebrafish Abcb4 and Abcb5 are functional transporters. The zebrafish Abcb4 protein was shown to transport all fluorescent P-gp substrates tested and was found to confer resistance to 90 P-gp substrates examined by high-throughput screening, while zebrafish Abcb5 conferred resistance to a narrower range of substrates. However, we also noted some differences in inhibitor specificity that were transporter- and substrate-dependent. Care must therefore be taken when evaluating the role of transporter inhibitors in modulating Abcb4 in order to increase brain penetration.

Although techniques for oral dosing of zebrafish have been developed [26], little is known in terms of the transporters expressed in the gastrointestinal tract. However, expression of *ABCB1* homologs has been described in other fish species [16], suggesting that this may also be true for the zebrafish. We found high reactivity with the C219 antibody in the gastrointestinal tract of the zebrafish and found that this was mainly due to high levels of *abcb4* expression. This is in agreement with Umans and Taylor who reported C219 staining in the intestinal epithelium of zebrafish larvae {Umans, 2012 #4892} and Lu et al. who found high levels of *abcb4* gene expression in the intestine of adult zebrafish [29]. The high levels of Abcb4 in the zebrafish gut appear to suggest that the zebrafish could be used in pre-clinical studies to determine the oral bioavailability of drugs. But, as mentioned above, the role that inhibitors might play to increase oral bioavailability would need to be scrutinized in light of the variable effects of P-gp inhibitors.

One lingering question with regard to transporters expressed at the zebrafish BBB and in the gut is the involvement of *ABCG2* homologs. Zebrafish have four direct *ABCG2* homologs— *abcg2a, abcg2b, abcg2c* and *abcg2d*—and little is known about their expression or substrate specificity. In one of the most comprehensive studies of these homologs, Kobayashi and colleagues found overexpression of zebrafish *abcg2a* and *abcg2c* in side-population cells isolated from the kidney, which is the main organ of hematopoiesis in the zebrafish [30]. Additionally, they reported expression of *abcg2a, abcg2b*, and *abcg2c* in the intestine [30]. Although cells transfected to overexpress zebrafish Abcg2a were found to transport Hoechst 33342, this appears to be the only known substrate of the zebrafish ABCG2 homologs [30]. Further work is needed to characterize these proteins and determine their potential role in the gut and BBB.

In summary, we find that expression of zebrafish Abcb4 at the zebrafish BBB phenocopies the role of human P-gp at the human BBB, suggesting that zebrafish Abcb4 could be considered a true homolog of P-gp. Although questions still remain in terms of other transporters such as ABCG2, the zebrafish may be a powerful tool to study the role of transporters at barrier sites.

## Acknowledgements

We appreciate the technical assistance of Christian Mustroph and the editorial assistance of George Leiman. This research was supported by the intramural programs at the National Cancer Institute and the National Center for Advancing Translational Sciences. The project has been funded in whole or in part with federal funds from the National Cancer Institute, National Institutes of Health, under Contract No. 75N91019D00024. The content of this publication does not necessarily reflect the views or policies of the Department of Health and Human Services, nor does mention of trade names, commercial products, or organizations imply endorsement by the U.S. Government. NCI-Frederick is accredited by AAALAC International and follows the Public Health Service Policy for the Care and Use of Laboratory Animals. Animal care was provided in accordance with the procedures outlined in the “Guide for Care and Use of Laboratory Animals” (National Research Council; 2011; National Academy Press; Washington, D.C.).

