## Supplementary material for "Characterization and tissue localization of zebrafish homologs of the human *ABCB1* multidrug transporter": Figure S1

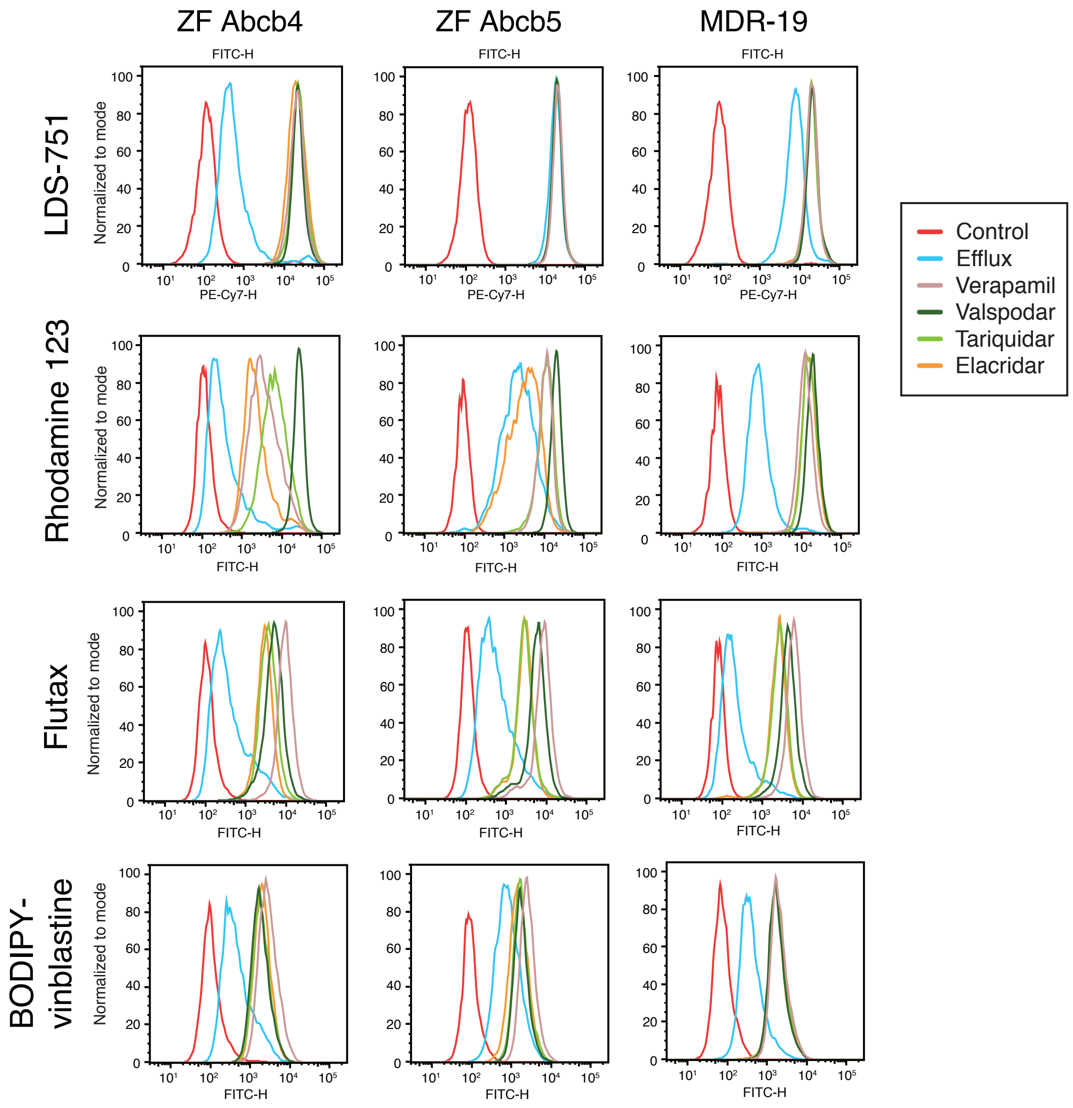


**Supplemental Figure S1**. Transport of fluorescent P-gp substrates by zebrafish Abcb4 and Abcb5. HEK293 cells transfected to express zebrafish Abcb4 (ZF Abcb4), zebrafish Abcb5 (ZF Abcb5) or human P-gp (MDR-19) were incubated in medium with (150 nM calcein-AM, 5 μM Flutax, 0.5 μM BODIPY prazosin, 250 nM BODIPY-vinblastine, 250 nM SYTO-13, 0.5 μM BODIPY FL-EDA, 0.5 μM TMRE or 0.5 μg/ml rhodamine 123) in the presence or absence of 10 μM elacridar, 10 μM tariquidar, 10 μM valspodar, or 100 μM verapamil for 30 min. The medium was removed and replaced with substrate-free medium in the presence or absence of inhibitor for an additional 1 h.
